# CROSS-GENERATIONAL IMPACT OF EPIGENETIC MALE INFLUENCE ON PHYSICAL ACTIVITY IN RAT

**DOI:** 10.1101/2022.03.21.485094

**Authors:** Sergey K. Sudakov, Natalia G. Bogdanova, Galina A. Nazarova

## Abstract

The aim of this work was to study whether epigenetic events at conception influence the formation of behavioral features found in adult rats. First generational inheritance of activity level, anxiety, and learning ability was studied. To separate genetic and non-genetic inheritance, mating of males and females with average motor activity was carried out in the presence males with high or low activity. Our results show that offspring of parents who mated in the presence of males with a high motor activity were significantly more active than offspring of parents that were paired in the presence of males with low activity. Anxiety level and learning ability were not inherited in this way. It is possible that the phenomenon we discovered is important for maintaining a certain level of activity of specific populations of animals. It counteracts natural selection, which should lead to a constant increase in the activity of animals.

## Introduction

Animals, including humans, have distinct individual characteristics which includes differences in the stability of visceral physiological functions as well as differences in the function of the brain and higher nervous activity. Individual behavioral characteristics can be inherited which directly affects fitness and influences population natural selection [1]. In addition to natural selection, we can use artificial selection to create populations of animals with given individual characteristics. Thus, in experimental physiology and pharmacology, selective strains of mice and rats with distinctive behavioral features can be used, for example, that differ in the level of motor activity [2,3], anxiety [4–6], or learning ability [7,8].

In addition to genetic mechanisms of inheritance, various epigenetic factors can also influence the formation of individual behavioral characteristics in offspring. These factors on a pregnant female include physical [9,10], chemical [11,12], and stressful influences [13,14]. Other environmental factors present in the postnatal and adolescent periods have also been reported to affect the formation of behavioral characteristics such as physical (15,16), chemical [17,18], and social influences [19–21], along with emotional stress [22]. Moreover, the behavior of animals can change even within the adult period [23] and the aging process itself can have a large impact on the behavior of animals and humans [24,25]. Thus, the initial genetic characteristics related to the formation of animal behavior are subject to significant epigenetic influences from the beginning of intrauterine development until death.

We hypothesized that it is possible to influence the behavior of future offspring at the time of conception. This influence can be imparted from the male during mating with a female, that is, in addition to the transfer of genetic material the male can also pass on epigenetic information relating to his behavioral characteristics.

The aim of this work was to study the still unexplored possibility of an epigenetic influence on the formation of behavioral features of an adult animal at the moment of its conception. The inheritance of high and low activity, high and low levels of anxiety, and high and low ability for spatial learning was studied in the first generation of rats. To distinguish between genetic and non-genetic inheritance, the rats were mated in the presence of several males with specific behavioral characteristics and these males were separated from the mating pair of animals by a latticed septum. A mated male and female had average values of motor activity, level of anxiety, or ability to spatial learning, but mating was done in the presence of living or dying males with high or low values for these indicators.

## Materials and Methods

### Animals

The experiments were carried out on Wistar rats weighing 210-240 g (male) and 170-200 g (female), were obtained from Stolbovaya Animal Clinic (Moscow, Russia). The animals were housed in ventilated cages Techniplast green line 1500U with natural corn bedding (Zolotoy Kot, Russia), 4-5 individuals in each, with free access to water and a standard combined food (3 kcal/g; Profgryzun, Russia) at a temperature of 21°C in the presence of lighting (90 Lux) from 20:00 to 08:00.

### Determination of inheritance of the level of motor activity

60 male and 20 female rats were individually placed in experimental chambers (Phenomaster, TSE, Germany) where horizontal locomotor activity was automatically measured for 60 min over intervals of 10 min. The experimental chambers were identical to the “home” cages in which the rats were housed in. The experiments were carried out from 11:00 to 15:00 in the absence of lighting.

Based on these results, 2 groups of males were formed with 15 animals in each group. The high activity (ACT) group had the highest horizontal motor activity (8758.13 ± 485.4 Units) and the passive (PAS) group had the lowest horizontal motor activity (1694.94 ± 194.8 Units). In addition, 6 males (MM) and 12 females (FM) were selected with average locomotor activity (4907.5 ± 455.4 and 4034.2 ± 425.2 Units, respectively).

The ACT group was randomized into 3 groups of 5 animals each. Three days later, one of 12 FM females in a state of estrus and an MM male, who began the mating process, were placed in the upper chamber of “two-story” cages for 5 min. 5 ACT males were housed on the bottom floor and separated from the top by a lattice partition. This process was repeated 3 times with the FM female in estrus along with one of the MM males and with each group of ACT males.

As a result, 13 female and 14 male offspring were obtained (groups ACTf1 and ACTm1, respectively). After 3 days, the experiment was repeated with PAS males according to the same scheme. Mating resulted in offspring of 18 females and 11 males (groups PASf1 and PASm1, respectively).

Three days after the first stage of the experiment, 5 ACT males were injected intraperitoneally with 500 mg/kg of sodium barbiturate and placed on the lower floor of the two-story cages; after 30 minutes, these animals were decapitated. 15 min before euthanizing the animals, an FM female in a state of estrus and an MM male were placed in the upper floor, which began the mating process. This process was repeated two more times in the presence of an FM female in a state of estrus with one of the MM males and with the other two groups of ACT males. As a result, 14 female and 16 male offspring were obtained (groups ACTf2 and ACTm2, respectively).

After the experiment with ACT males was complete, the experiment was repeated with PAS males in the same way. Mating resulted in 18 female and 11 male offspring (groups PASf1 and PASm1, respectively) and 17 females and 12 males (groups PASf2 and PASm2, respectively).

At the age of two months, the level of horizontal motor activity was measured in the ACT and PAS offspring according to the method described above. At the age of 5 months, the study of motor activity was repeated.

### Determination of inheritance of the level of anxiety

In the second series of experiments, 60 male and 20 female rats were studied in two successive experimental tests: “open field” (OF) and “elevated plus maze” (EPM). The tests were carried out at intervals of 7 days.

The OF test was a circular arena 98 cm in diameter, surrounded by walls 31 cm high ([OpenScience](https://translate.google.com/translate?hl=ru&prev=_t&sl=ru&tl=en&u=http://www.openscience.ru/index.php?page=ts&item=001), Russia, model TS0501-R). The arena was divided by lines into 7 central and 12 peripheral sectors. The arena was uniformly illuminated by a medical lamp (illumination of the arena surface = 450 Lux). For the OF test, a rat was placed in the center of the arena and horizontal activity (total number of crossed squares) was observed for five min. The animal was returned to its home cage after the test.

The EPM test (The Elevated Plus Maze, Columbus Instruments, USA) uses a cruciform platform with four arms (length = 50 cm, width = 15 cm). Two opposite arms of the maze had high, opaque walls, whereas the other two were open. The height of the walls of the closed arms were 43 cm. The maze was raised to a height of 75 cm. The central part of the EPM was 15 cm^2^. Illumination of the surface of the maze was 90 Lux. For this test, a rat was placed for 5 min in the center of the EPM platform and the individual anxiety of the animals was determined by the time spent on the open arms of the EPM.

According to the method described earlier [26], 16 low-anxiety (LA) males were selected with the highest intensity of motor activity in the OF test (22.06 ± 2.43 crossings) and the largest amount of time spent in EPM open arms (40.12 ± 6.05 s), and 16 high-anxiety (HA) males were selected with the lowest locomotor activity in the OF test (8.18 ± 1.43 intersections) and lowest time spent in the EPM open arms (4.18 ± 1.87 s). In addition, 4 males (MM) and 4 females (FM) were selected with average values (15.22 ± 2.1 crosses and 22.5 ± 4.95 s).

Three days after the end of testing, HA males were divided into 2 groups of 8 animals each. Eight HA males of the first group were intraperitoneally injected with 500 mg/kg of sodium barbiturate and placed on the lower floor of the cages, and 30 min later the animals were decapitated. Fifteen min before euthanizing the animals, an FM female in a state of estrus and an MM male were placed in the upper floor cage, which began the mating process. This process was repeated once more with the FM female in estrus with one of the MM males and 8 HA males of the second group.

As a result, 9 female and 13 male (HAf and HAm) offspring were obtained.

Three days later, an experiment with the LA group was carried out according to a similar scheme. As a result of these experiments, 9 female and 14 male offspring were obtained (groups LAf and Lam, respectively).

At the age of two months, the behavior of the animals was investigated using the OF and EPM tests, and at the age of 5 months the animals were re-examined with these tests.

### Determination of the inheritance of the learnability

In a first series of experiments, 44 male rats were first trained in a Morris water maze test ([OpenScience](https://translate.google.com/translate?hl=ru&prev=_t&sl=ru&tl=en&u=http://www.openscience.ru/index.php?page=ts&item=001), Russia, model TS1004-M2G). The maze was a round pool 1.8 m in diameter and 0.6 m deep, half filled with water with a temperature of 24°C. Lighting lamps were positioned in such a way as to prevent reflections on the water (30 Lux). The pool was divided into 4 sectors and a platform with a diameter of 12 cm was placed in one sector (called “target”) and submerged 1.5 cm below the water level. Four visual cues measuring 50 cm^2^ of various shapes were located 80 cm from the center of the maze and served as an external reference point. The rat was placed in the pool at the edge opposite to the platform and at a distance of 150 cm from it. The animal, swimming in a randomly chosen direction, would eventually find a platform and climb onto it. If the rat did not find the platform within 60 s, it was forcibly placed onto it.

On the day of training, the animals were presented with 4 sessions. Each animal was given 60 s to find the platform. If the rat on the first day of the experiment did not learn to find the platform on its own, it was removed from further training. The rest of the animals were trained for 4 days, 4 sessions per day. After training, two groups of animals were formed, which were used in further experiments. The first group (L) included 20 trained animals and at the end of the fourth day of training the latency period for reaching the platform was no more than 10 s. The second group (NL) consisted of 24 rats, which on the first day of 4 sessions never found an underwater platform on their own.

10 L males were injected intraperitoneally with 500 mg/kg of sodium barbiturate and placed in the lower floor cage, and 30 min later the animals were decapitated. Fifteen min before this, an FM female in a state of estrus and an MM male were placed in the upper floor cage, which began the mating process. This process was repeated once more with the FM female in estrus with one of the MM males and the remaining 10 L males. For mating MM and FM, we used animals with an average level of locomotor activity, as determined in the Phenomaster described above.

As a result of these experiments, 9 female and 9 male offspring were obtained (groups Lf and Lm, respectively). Three days later, the above procedure was repeated using NL rats. As a result of mating, 11 female and 11 male offspring were obtained. At the age of two months, the animals of the Lm and Lf groups and the NLm and NLf groups were tested in the Morris maze: the rats were placed in the maze 4 times a day with an interval of 1 min for 4 days. The latency period for reaching the underwater platform was measured. The animals were re-tested at the age of 5 months.

### Statistics

Statistics were performed using the statistical software Statistica 10.0. The data obtained in each experimental subgroup was assessed for normality using the Shapiro-Wilk and Kolmogorov-Smirnov tests. Since the test showed the absence of a normal distribution, the calculations were carried out by the nonparametric Mann-Whitney U test (Phenomaster, OF, and EPM data) and Van der Waerden One-Way Analysis (Morris maze data). Results in tables and figures are presented as the mean ± SD. The results were assessed as significant at p < 0.05.

### Animal welfare

The protocols and procedures for this study were ethically reviewed and approved by the Animal Care and Use Committee of the P.K. Anokhin Research Institute of Normal Physiology (Permission number 342) and conform to Directive 2010/63/EU.

### Groups of rats

**ACT -** active rats

**PAS -** passive rats

**ACTf1, ACTm1 -** female and male offspring of rats mated in the presence of live active rats

**PASf1, PASm1 - -** female and male offspring of rats mated in the presence of live passive rats

**ACTf2, ACTm2 -** female and male offspring of rats mated in the presence of dying active rats

**PASf2, PASm2 -** female and male offspring of rats mated in the presence of dying passive rats

**HA -** highly anxious rats

**LA -** low anxious rats

**HAf, Ham -** female and male offspring of rats mated in the presence of dying highly anxious rats

**LAf, LAm -** female and male offspring of rats mated in the presence of dying low anxious rats

**L -** learned rats

**NL -** non-learned rats

**Lf, Lm -** female and male offspring of rats mated in the presence of dying learned rats

**NLf, NLm -** female and male offspring of rats mated in the presence of dying non-learned rats

## Results

### Determination of inheritance for the level of motor activity

The offspring of PAS rats at the age of 2 months had significantly lower motor activity than the offspring of ACT rats. Moreover, the differences were between the groups of offspring obtained in the first experiment in males ACTm1 and PASm1 (U=28.00000, *p*=0.006217, Z=2.655127) and females ACTf1 and PASf1 (U=60.50000, *p*=0.022087, Z=2.241794), and in the second experiment in males ACTm2 and PASm2 (U=26.00000, *p*=0.000670, Z=3.266457) and females ACTf2 and PASf2 (U=20.00000, *p*=0.000419, Z=3.989268). The differences in the second experiment, in the presence of dying animals during conception, were more significant than in the first experiment, when live animals were present at conception (Fig.s 1,2). Thus, in ACTm2 males and ACTf2 females, the level of motor activity was significantly higher than in ACTm1 males and ACTf1 females (U=31.00000, *p*=0.000417, Z=-3.34643; U=11.00000, *p*=0.00035469, Z=3.822814, respectively). The PASf2 females had a significantly lower level of motor activity than the PASf1 females (U=31.00000, *p*=0.027473, Z=2.887306). There were no differences in the PASm1 and PASm2 groups.

**Figure 1.**
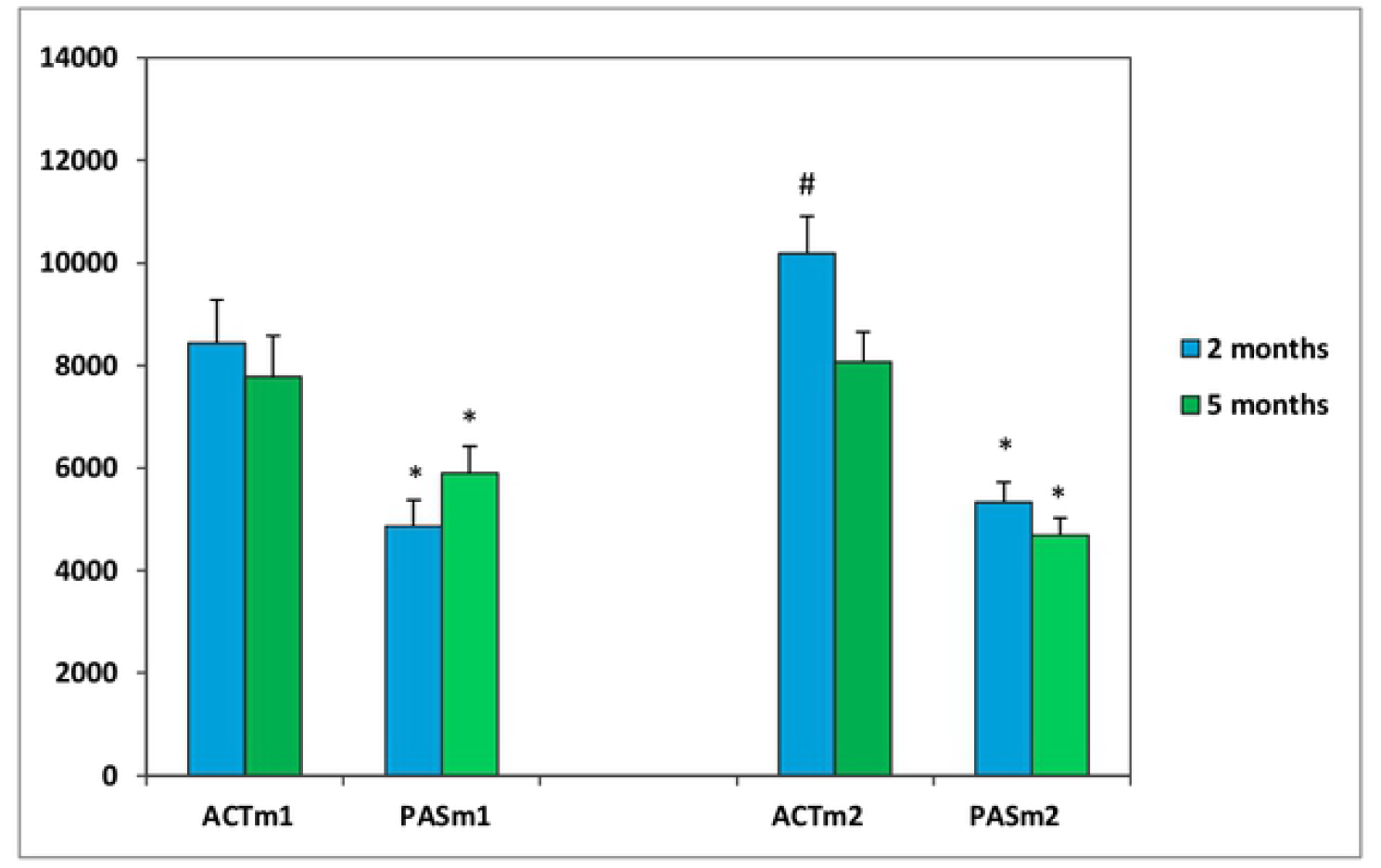
Total horizontal motor activity of male rats for 60 min (conventional units). * *p*<0.05 between ACTm and PASm; ^#^ *p*<0.05 between ACT1 and ACT2.

**Figure 2.**
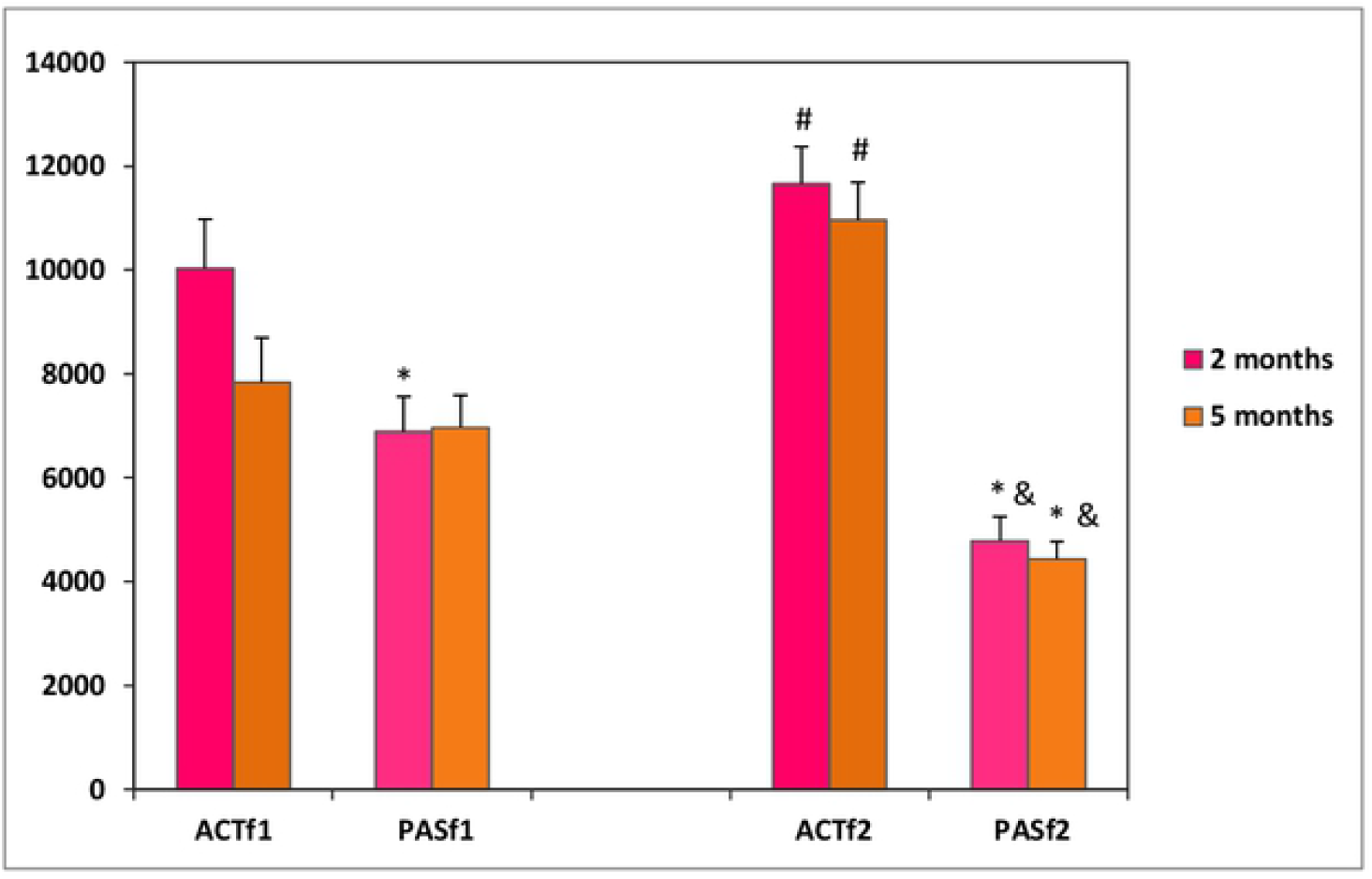
Total horizontal motor activity of female rats for 60 min (conventional units). * *p*<0.05 between ACTf and PASf; ^#^ *p*<0.05 between ACT1 and ACT2; ^&^ *p*<0.05 between PAS1 and PAS2.

At the age of 5 months, significant differences in the level of motor activity between the male PAS and ACT offspring remained both between ACTm1 and PASm1 (U=39.00000, *p*=0.038425, Z=2.052933) and between ACTm2 and PASm2 (U=30.00000, *p*=0.000315, Z=2.922814) (Fig. 1). In females, significant differences between the ACTf1 and PASf1 groups disappeared by 5 months. However, there was a significant difference in motor activity between the ACTf2 and PASf2 groups (U=17.00000, *p*=0.000781, Z=3.644271) (Fig. 2). In 5-month-old PASf2 females, as well as in 2-month-old females, a significantly lower level of motor activity was observed than in PASf1 females (U=51.00000, *p*=0.000467, Z=3.333497).

### Determination of inheritance for the level of anxiety

HA offspring at the age of 2 months had a significantly shorter time spent in the center of the OP test than the LA offspring. These differences were significant both between HAm and LAm males (U=50.00, *p*=0.048173, Z=1.9653) and between HAf and LAf females (U=17.00, *p*=0.042261, Z=-2.030949) (Fig. 3). At the age of 5 months, significant differences in this parameter in the offspring of HA and LA disappeared. Significant differences in motor activity were also observed between these groups of animals. Both HAm (U=37.50, *p*=0.007796, Z=2.580566) and HAf (U=6.50, *p*=0.001234, Z=2.958122) at the age of 2 months had lower motor activity in the OF test than LAm and LAf (Fig. 4). By the 5th month of life, these differences disappeared.

**Figure 3.**
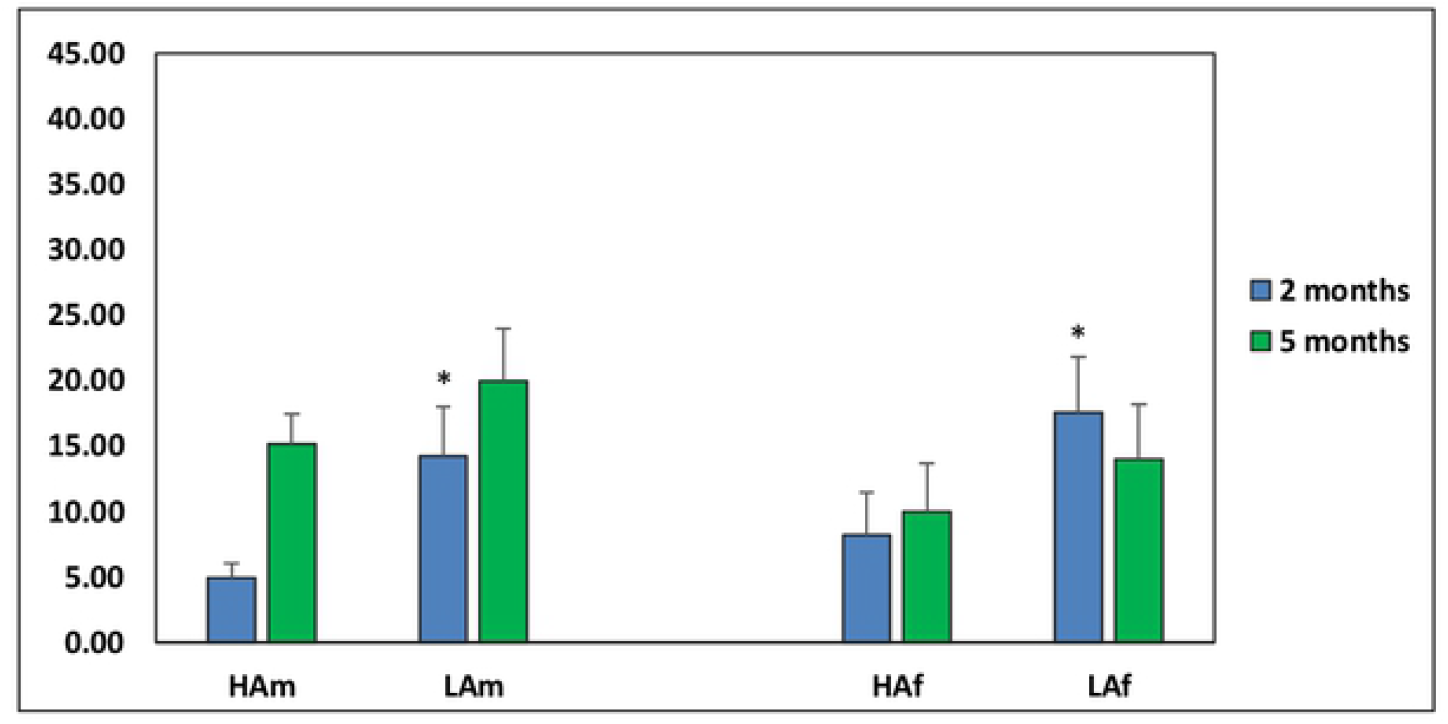
Time spent at the open field center. * p <0.05 between HAm and LAm and HAf and LAf 2-month old.

**Figure 4.**
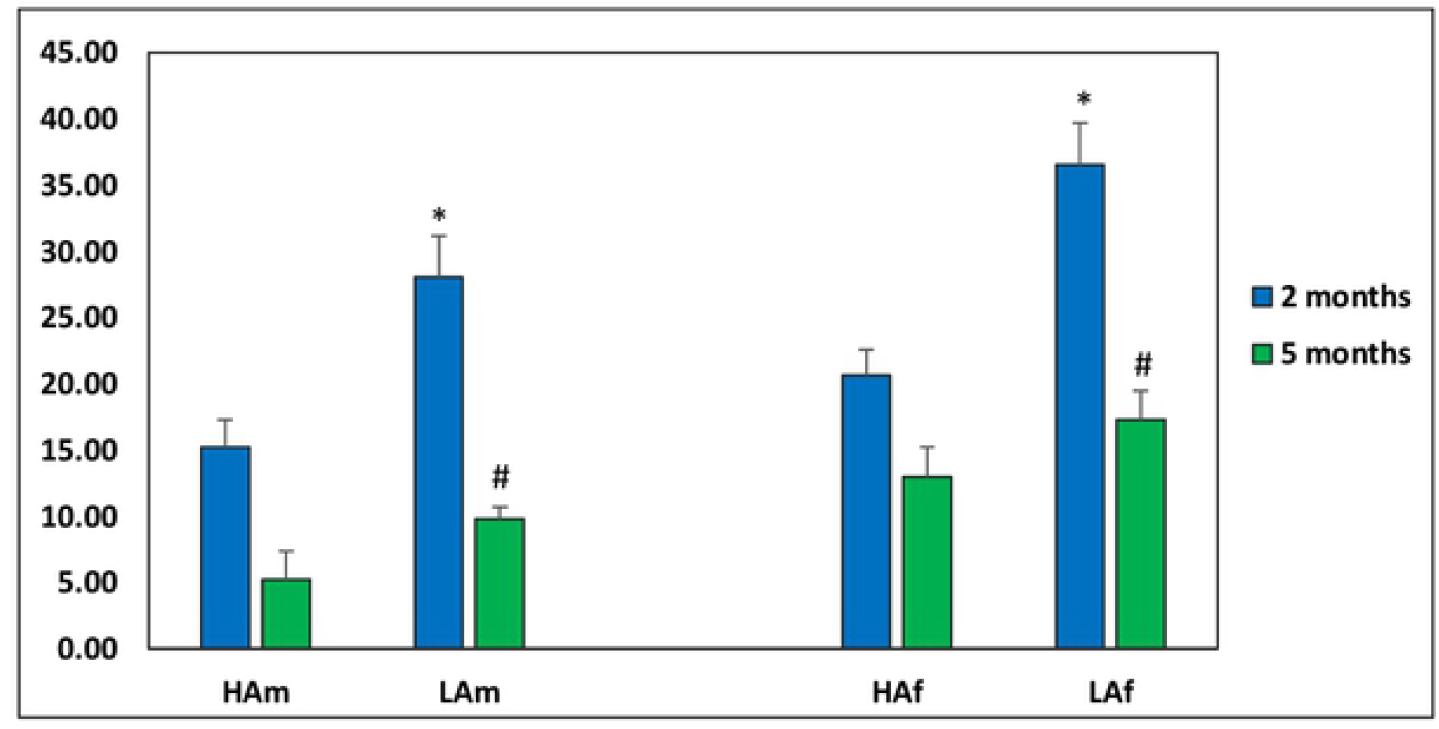
Number of intersections in an open field test. * p <0.05 between HAm and LAm 2 month old and HAf and LAf 2 month old; ^#^ p <0.05 between LAm 2- and 5-month and LAf 2- and 5-month old.

In addition, both in LAf (U=4.00, *p*=0.000494, Z=3.178878) and in LAm (U=10.50, *p*=0.000007, Z=3.997448) animals at the age of 5 months had a significantly decreased level of motor activity. At the same time, HA animals did not show a change in the level of motor activity in the OF test from the 2nd to 5th month of age (Fig. 4).

There were no significant differences between the groups in the time spent on the open arms of the EPM test (Fig. 5). Both LAf (U=3.00, *p*=0.000288, Z=3.2671) and Lam (U=36.50, *p*=0.006618, Z=2.620413) animals at 2 months of life were significantly more active than HA rats. In addition, the motor activity of LAm (U=28.00, *p*=0.000794, Z=3.193364) and LAf (U=12.00, *p*=0.010613, Z=2.472460) animals was significantly decreased from 2 to 5 months of age (Fig. 6).

**Figure 5.**
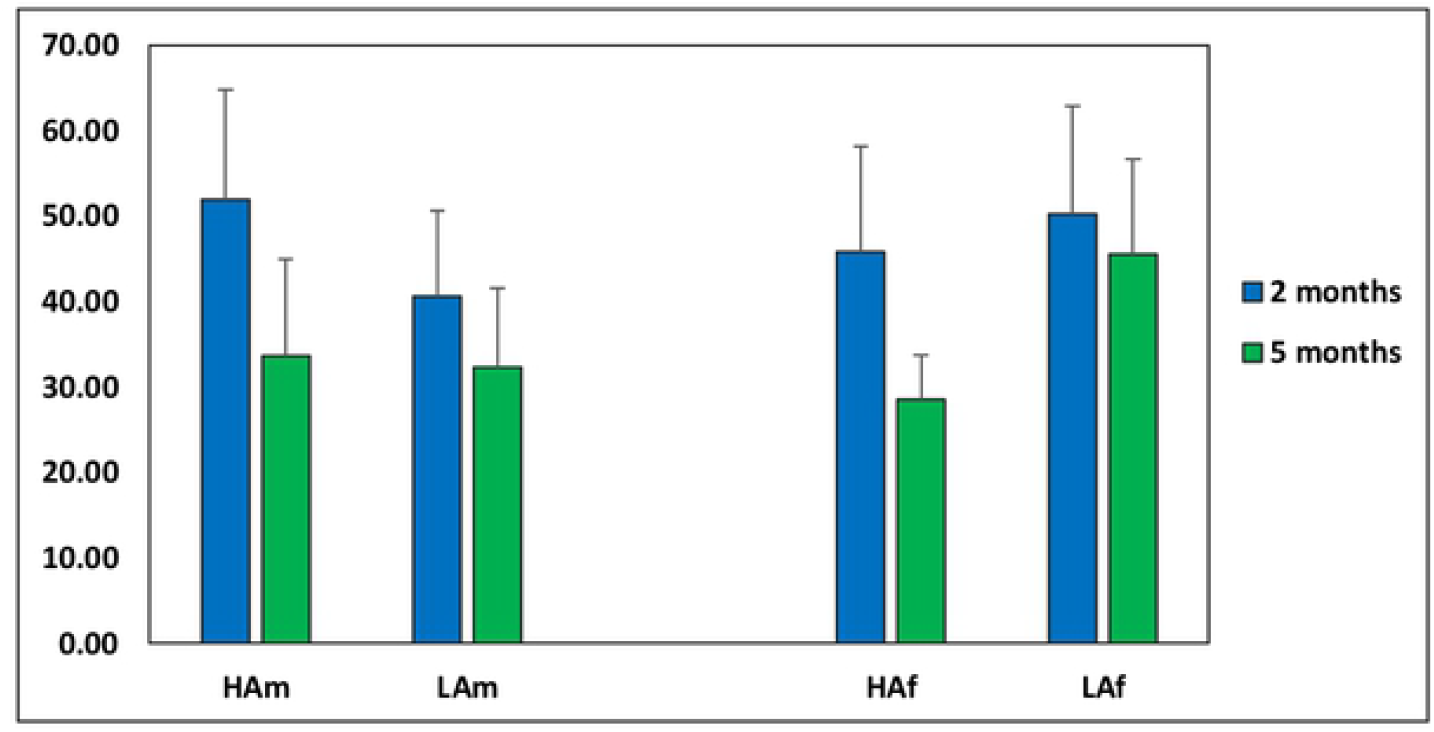
Time spent on the elevated plus maze open arms.

**Figure 6.**
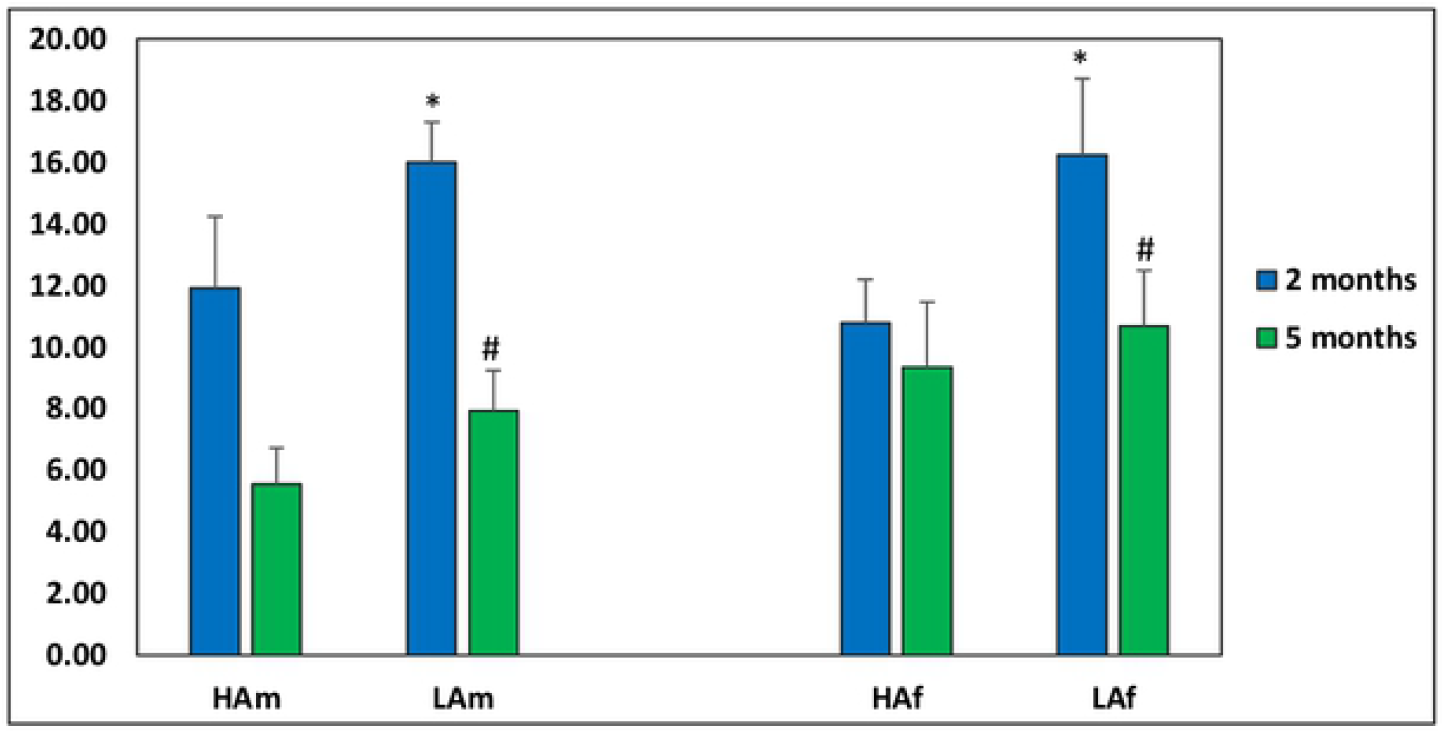
The number of intersections crossed in the elevated plus maze test. *\* p*<0.05 between HAm and LAm and HAf and LAf 2 month old; ^#^ *p*<0.05 between LAm 2- and 5-month and LAf 2-and 5-month old.

### Determination of inheritance of learnability

The animals conceived in presence of the L group differed from the animals conceived in the presence of the NL group in the speed of reaching the underwater platform in the Morris maze. Thus, in the first days of the experiment, Lm rats had a longer latency period for reaching the platform than rats of the NLm group; these changes were significant and most pronounced on the third day of testing (Pr > Chi-Square - 0.0095; 0.0292). On the fourth day of the experiment, the differences completely disappeared (Fig. 7a). Lf rats practically did not differ from NLf rats in the latency period of reaching the underwater platform in the first two days of the experiment. However, on the third day, and especially on the fourth day, the females of the Lf group reached the underwater platform significantly faster than the females of the NLf group (Pr > Chi-Square - 0.0085; 0.011) (Fig. 7b).

**Figure 7.**
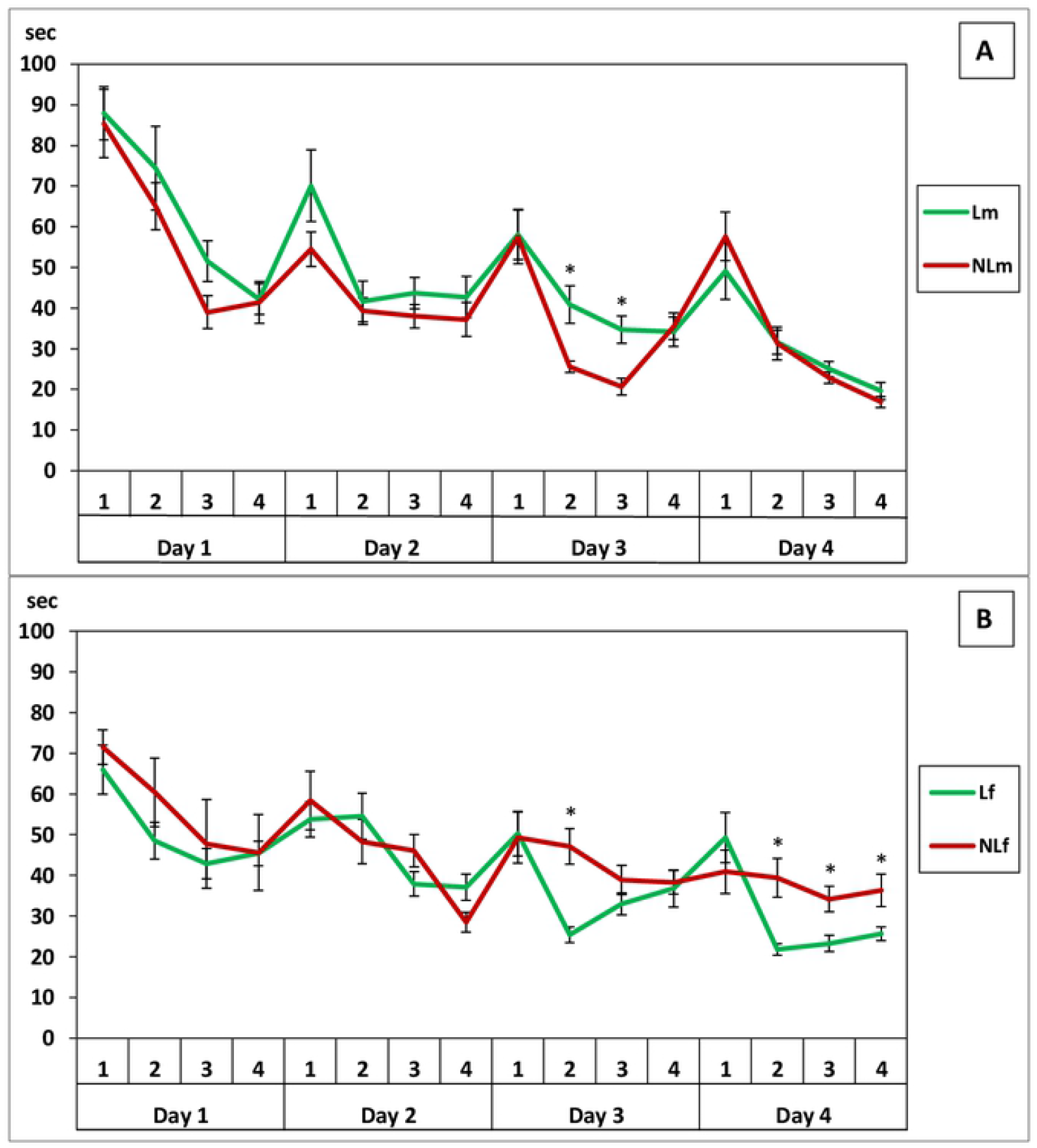
The latency period of reaching the underwater platform (s) (7a - 2-month male rats; 7b - 2-month female rats). * *p*<0.05 between L and NL rats.

When tested again at the age of 5 months, it was found that on the first day Lm males had a longer latency period for reaching the platform, and on the fourth day Lm males reached the platform significantly faster than NL males (Pr > Chi-Square - 0.0004; 0.012) (Fig. 8a). The females retained the differences between the groups observed during the first testing: on the first two days, there were no differences, and on the third day, and especially on the fourth day, Lf rats reached the underwater platform much faster (Pr > Chi-Square - 0.03; 0.0015) (Fig. 8b).

**Figure 8.**
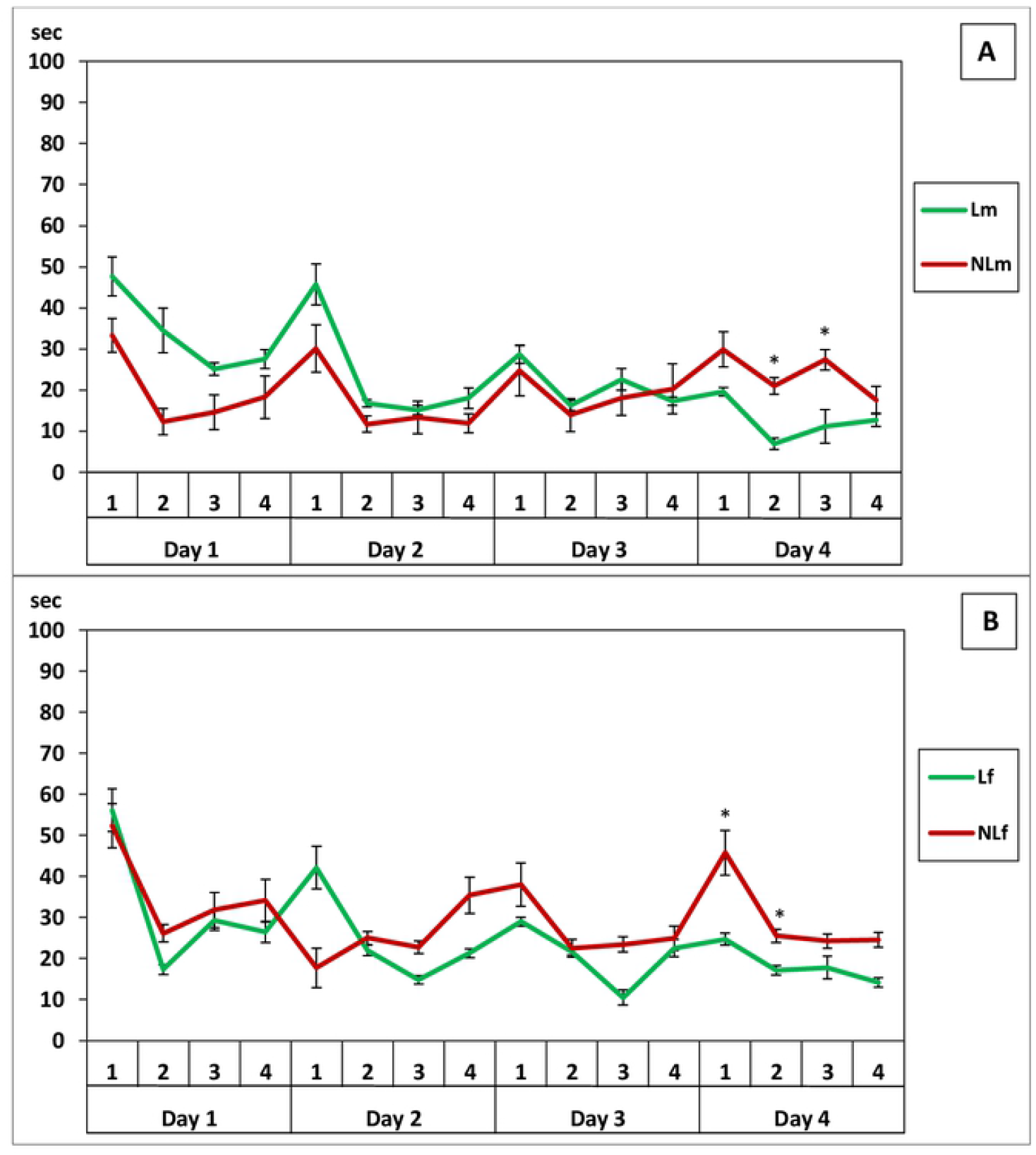
The latency period of reaching the underwater platform (s) (8a - 5-month male rats; 8b - 5-month female rats). * *p*<0.05 between L and NL rats.

## Discussion

The results of this work show that even at the moment of conception exertion of epigenetic influences, which have a significant impact on the behavior of future offspring, are possible. Our experiments made it possible to separate the genetic component derived from the biological father and epigenetic information which was passed on to the female from mating in the presence of living or dying males. Perhaps in nature both of these components come from the same male participating in mating. Thus, independently of static genetic information, future offspring will have the same level of activity that is found in the father during mating and the male can have different levels of physical activity at different times of the childbearing period [23].

We found that this epigenetic inheritance is only related to the level of activity of the fathers and offspring. Anxiety and spatial learning were not similarly inherited. The identified changes appear to be related to the level of activity. Thus, males, donors of epigenetic information, with a higher level of activity, may more often visit the center of the open field and open arms of the EPM test [27]. Also, active animals can find an underwater platform in the Morris maze faster [28]. Thus, we believe that epigenetic inheritance is influenced by the level of motor activity from the male, which either participates in the mating process or is nearby.

Most likely, the epigenetic influence from the male passes to the female participating in the mating process. This effect is quite long-term and can have an impact on the bearing of offspring and their upbringing. The nature of this epigenetic effect is still unclear. Two hypotheses can be suggested. The first, and most probable, is that the impact is mediated by odors (pheromones). For example, primer pheromone effects found in the house mouse were associated with a change in its hormonal status under the influence of male pheromones [29–34]. In addition, presentation of the male pheromone before mating has been found to enhance mammary development during pregnancy, which is reflected in the development of the mammary glands in the postpartum period during breastfeeding [35]. It was also shown that in offspring obtained from females receiving male pheromone, there is an increase in the expression of polysialyltransferases in the brain which have been shown to be involved in brain development [36]. Subsequently, such offspring showed a greater ability for spatial learning (35). In our experiments, we found that the presence of dying males in the immediate vicinity of mating had the greatest epigenetic effect on the future level of motor activity in offspring. This effect has yet to be explained. Perhaps in the process of dying the male secretes more pheromones. However, we could not find such data in the literature to support this.

The second hypothesis is that a magnetic, electromagnetic, or electrical force can be transferred from male to female. This concept has not been studied in depth, although there is data on the effect of electromagnetic radiation immediately after fertilization on further fetal development in mice [37]. Additionally, there is evidence that during death, rats exhibit increased electrical activity (a ‘death wave’) [38], which theoretically can affect the body in the vicinity of the dying subject.

The results of our experiments show that epigenetic inheritance of a particular motor activity is carried out by both male and female offspring. The differences between ACTm and PASm are similar to those between ACTf and PASf. This may indicate that the epigenetic influence is carried out regardless of the sex of the offspring. However, there were some differences between males and females. Possibly, these differences could be related to the fact that testing of females in Phenomaster, OF, and EPM tests could have occurred in different hormonal states of females, which is known to affect behavior in this test [39,40].

There are certain limitations of this study that require further experimentation. First, there was a small number of litters in each group. Despite the rather significant number of offspring obtained from mating of parents in the presence of ACT or PAS rats, there were only three litters in each group. Nevertheless, the statistical analysis showed high reliability in the differences between groups, which would unlikely arise from random genetic characteristics of a small number of parents. However, increasing amounts and litters in each treatment group would be appropriate. Second, the nature of the epigenetic effect is not clear. Future experiments in this direction will be aimed at eliminating the possibility of transferring odors from ACT or PAS animals or, conversely, presenting their pheromones to mating animals. However, this would investigate the epigenetic influence only from males. In further experiments, females will be used as possible donors of epigenetic effects on the future behavior of their offspring.

It is possible that the phenomenon we discovered is important for maintaining a certain level of activity of specific populations of animals. It counteracts natural selection, which should lead to a constant increase in the activity of animals.

## Acknowledgments

We are grateful to E.A. Alekseeva for technical assistance. We thank A.Yu. Abramova and V.P. Leonov for help in statistical analysis.

Editorial assistance, in the form of language editing and correction, was provided by XpertScientific Editing and Consulting Services.

